# Tocotrienol decreases β-amyloid mediated toxicity in *Caenorhabditis elegans* model of Alzheimer’s disease

**DOI:** 10.1101/2020.04.13.040105

**Authors:** Chee Wah Yuen, Mardani Abdul Halim, Vikneswaran Murugaiyah, Nazalan Najimudin, Ghows Azzam

## Abstract

Alzheimer’s disease (AD) is a neurological disease caused by the accumulation of extracellular senile plaques consisting of β-amyloid peptide (Aβ) in the brain. A transgenic *Caenorhabditis elegans* which demonstrated paralysis due to the expression of human beta amyloid Aβ_42_ gene was used to study the anti-paralysis effect of mixed tocotrienols. The content of the mixed tocotrienols were 12.1% α-, 2.7% β-, 18.6% γ-, and 8.1% δ-tocotrienols. Mixed tocotrienols significantly delayed the Aβ-induced paralysis in the transgenic nematode and exhibited anti-oxidant properties towards Aβ-generated oxidative stress. The mixture also presented potent inhibitory activities against Aβ aggregation with an IC_50_ value of 600 ng/ml. It is concluded that mixed tocotrienols could potentially serve as a new therapeutic candidate for AD.

## 1.0 Introduction

Alzheimer’s disease (AD) is a progressive neurodegenerative disorder that affects the old age population resulting in the loss in memory loss and decrease muscle strength. The occurrence of this disease is due to the presence of neurofibrillary tangles and senile plaques in the brain (1). Aβ is a protein that aggregates to form fibrils in the human brain and is expressed by a gene in chromosome 21 in humans (2). The manifestation of AD occurs when Aβ amyloid plaques start to materialize in the basal part of the isocortex and subsequently spread to the rest of the isocortex due to progression of AD. At the later stage of AD, the amyloid plaques have covered almost the brain with the highest amyloid plaque density found in the isocortex (3).

Tocotrienols are a group of chemicals forming part of the vitamin E family and normally found in the seed endosperm of many monocotyledons like palm, wheat, rice and barley and some dicotyledons like tobacco (4). Palm oil is one of the natural sources of tocotrienols and approximately 800 mg tocotrienols per kg crude oil can be extracted from it. Mixed tocotrienols has been reported to have potential neuroprotective properties in preventing Aβ aggregation. Inhibition of Aβ_42_ fibrillogenesis were observed when Aβ_42_ peptide was exposed to mixed tocotrienols with the composition 19.6% α-, 2.4% β-, 25.5% γ- and 7.5% δ-tocotrienols. Aβ_40_ and Aβ_42_ oligomer formations were also suppressed by the mixed tocotrienols. In addition, mixture also reduced Aβ deposition in the brain and significantly improved the cognitive functions of AβPPswe/PS1dE9 double transgenic mice (5).

Transgenic *Caenorhabditis elegans* containing the human Aβ_42_ gene have been used as a model in various studies investigating the effect of natural products on Alzheimer’s disease (6). The short life span of the nematode, easy maintenance and its concomitant progressive paralysis phenotype makes it favorable as a model. In addition, it has the ability to develop muscle-associated deposits reactive to amyloid-specific dyes. It is also experiencing Aβ_42_-induced oxidative stress (7). Thus far, no study on the effects of tocotrienol on the transgenic *C. elegans* has ever documented. Hence, the objective of this paper is to demonstrate the efficacies of mixed tocotrienols extracted from *Elaeis guineensis* (oil palm) in attenuating Aβ-mediated toxicity in transgenic nematodes that carry the Aβ_42_ protein gene.

## 2.0 Methods and Materials

### 2.1 *C. elegans* strains and Maintenance

Transgenic *C. elegans* strain GMC101(dvIs100 [unc-54p::A-beta-1-42::unc-54 3’-UTR + mtl-2p::GFP), *C. elegans* strain CL2122 (dvIs15 [(pPD30.38) unc-54(vector) + (pCL26) mtl-2::GFP]) and *E. coli* OP50 strain were kindly provided by the Caenorhabditis Genetics Center (CGC), University of Minnesota (Minneapolis, MN, USA). The *C. elegans* strain were maintained at 16 °C on nematode growth medium (NGM) seeded with *E. coli* OP50 bacteria. GMC101 is a transgenic strain that expressed the human Aβ_42_ gene while CL2122 does not have the human Aβ_42_ gene.

### 2.2 *C. elegans* paralysis assays

Paralysis assays were performed as described by McColl *et al*. (8) with a slight modification. All populations were cultured at 16°C on 60 mm NGM plates with 250 μl of overnight *E. coli* OP50 bacterial culture pre-spread on the plates. After a 4-hour egg-laying period on NGM plates at 16°C, either in the absence or presence of the compounds, the *C. elegans* populations were developmentally synchronized by shifting the incubation temperature to 25°C at 72 hours post-egg lay. Time zero was defined at this point of temperature shift. The body movement of the nematodes were assessed over time by scoring as “paralysed” if they failed to show complete full body movement, either spontaneously or touch-provoked. The proportion of individuals that were not paralysed was calculated. Three independent experiments were performed in all studies. A range of different concentrations of mixed tocotrienols (0.25%-1.00%) from the oil palm *Elaeis guineensis* (from ExcelVite Pty Ltd) were used to treat the nematodes.

### 2.3 Measurement of reactive oxygen species (ROS) in C. elegans

The levels of ROS were measured *in vivo* by using the 2,7-dichlorofuorescein diacetate method described by Gutierrez-Zepeda *et al.* (7) with a modification in which the worms were incubated at 37°C between reads to simulate human body temperature. Readings were taken every 20 minutes. To initiate Aβ-induced paralysis, the live worms were shifted to an incubator set at 25°C. Approximately 100 worms were harvested at 32 hrs after temperature shift using M9 buffer. The worms were resuspended in M9 buffer with 100 μM DCFH-DA (Sigma-Aldrich, Missouri, USA). They were then transferred into the wells of a 96-well non-binding black microplate (Greiner Bio-One) and the fluorescence was read every 20 minutes at an excitation wavelength of 485nm and an emission wavelength of 520nm using a fluorescence microplate reader (EnVision 2104 Multilabel Reader (Perkin Elmer, MA, USA).

### 2.4 Immuno-dot blot assay of Aβ

The levels of Aβ proteins were analysed using the immuno-dot blot analysis. Samples were then extracted in 3 volumes of urea buffer (7 M urea, 2 M Thiourea, 4% w/v CHAPS, 1.5% w/v dithiothreitol and 50 mM Tris pH 8.0), disrupted via sonication and subsequently centrifuged at 13,000 x*g* for 10 min. Total proteins were spotted (10 μg) onto nitrocellulose membranes (Merck Millipore, Darmstadt, Germany) and left to dry. The membranes were incubated with a blocking solution [skim milk (Sunlac) with 0.1% tween-20-phosphate buffered saline] for one hour. This was followed by an incubation period with Anti-Aβ mouse monoclonal antibody 6E10 (1:1000, epitope: Aβ_3–8,_ Biolegend, San Diego) and peroxidase-conjugated anti-mouse IgG (1:1000, Bio-Rad, California) which acted as the primary and secondary antibodies, respectively. The spots were viewed using the colorimetric method available in Opti4CN™ Substrate Kit (Bio-Rad, California).

### 2.5 Aβ Aggregation assay

The procedure was performed as described by the manufacturer’s instructions (SensoLyte ® Thioflavin T Beta Amyloid (1-42) Aggregation Kit (Anaspec)). Approximately 10 μl of 2mM ThT, 85μL of Aβ_42_ peptide and 5μL of the bioactive compounds at selected concentrations were added into each well. The plates were read every 10 minutes on a fluorescence microplate reader (EnVision 2104 Multilabel Reader (Perkin Elmer, MA, USA) at an excitation wavelength of 440nm with an emission wavelength of 484nm.

## 3.0 Results

Paralysis of the worms were significantly delayed when worms were treated at 0.5 (50 µg/ml) %, 0.75% (75 µg/ml) and 1% (100 µg/ml) of mixed tocotrienols (p<0.05) (Figure 1 A and B). The lower percentage of tocotrienols (0.125% - 0.25%) did not significantly reduce paralysis.

**Figure 1:**
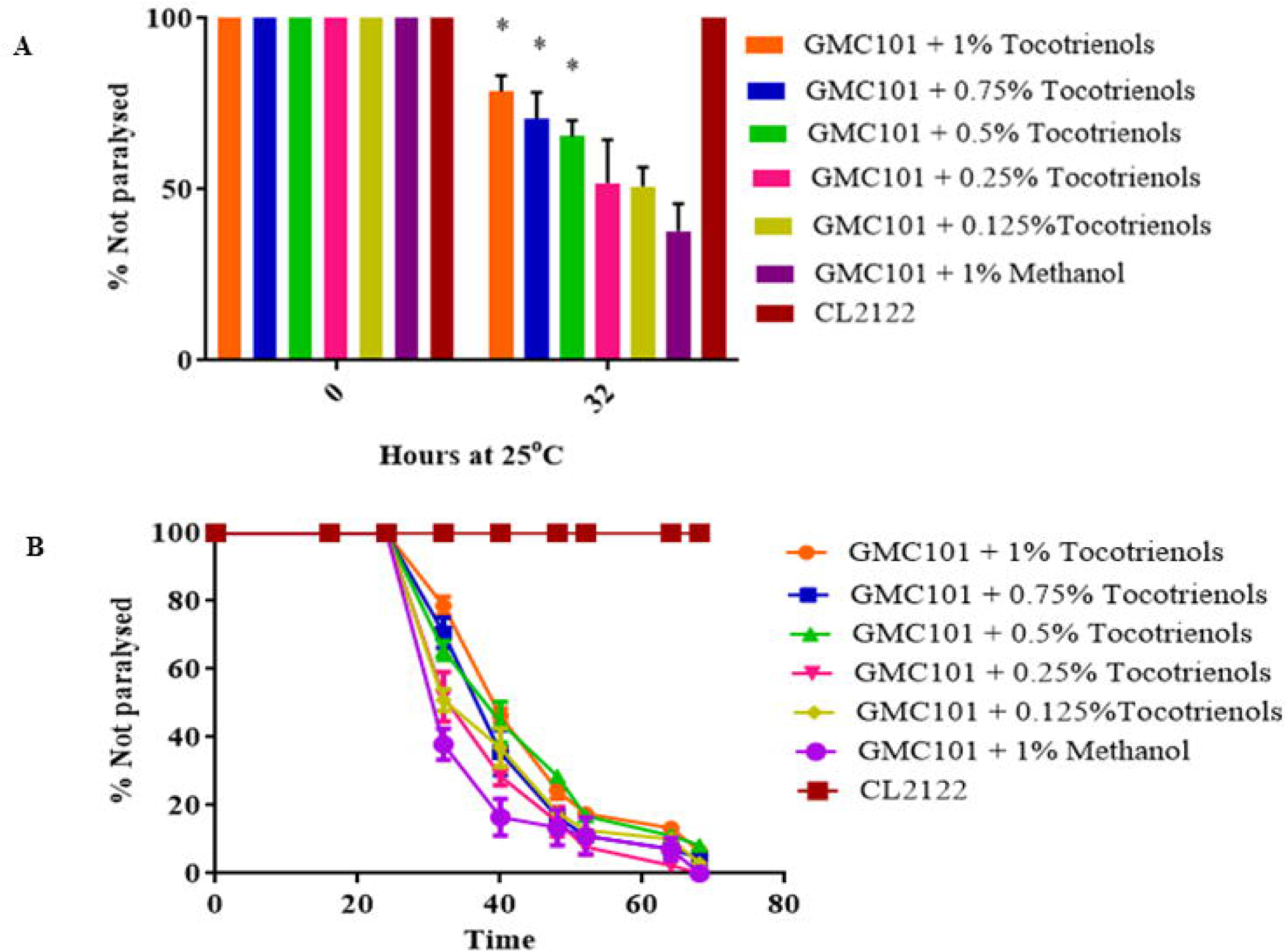
Effects of mixed tocotrienols on the paralysis of the worms. **A.** Graph showing anti-paralysis effects of transgenic *C. elegans* of different concentrations of mixed tocotrienols. **B.** Comparison of transgenic *C. elegans* strain GMC101 treated at various concentrations of percentages of tocotrienols at 0 and 32 hours. Paralysis of the worms were significantly delayed at 0.5%, 0.75% and 1% tocotrienols (p<0.05) at 32^nd^ hour after upshift from 16°C to 25 °C.

Aggregation assay using thioflavin T dye was used to measure Aβ_42_ aggregation in the presence of mixed tocotrienols. Concentrations of 200 ng/ml, 400 ng/ml, 600 ng/ml and 800 ng/ml of mixed tocotrienols were used to test their ability to produce anti-aggregation properties towards Aβ_42_ (Figure 2 A).

**Figure 2.**
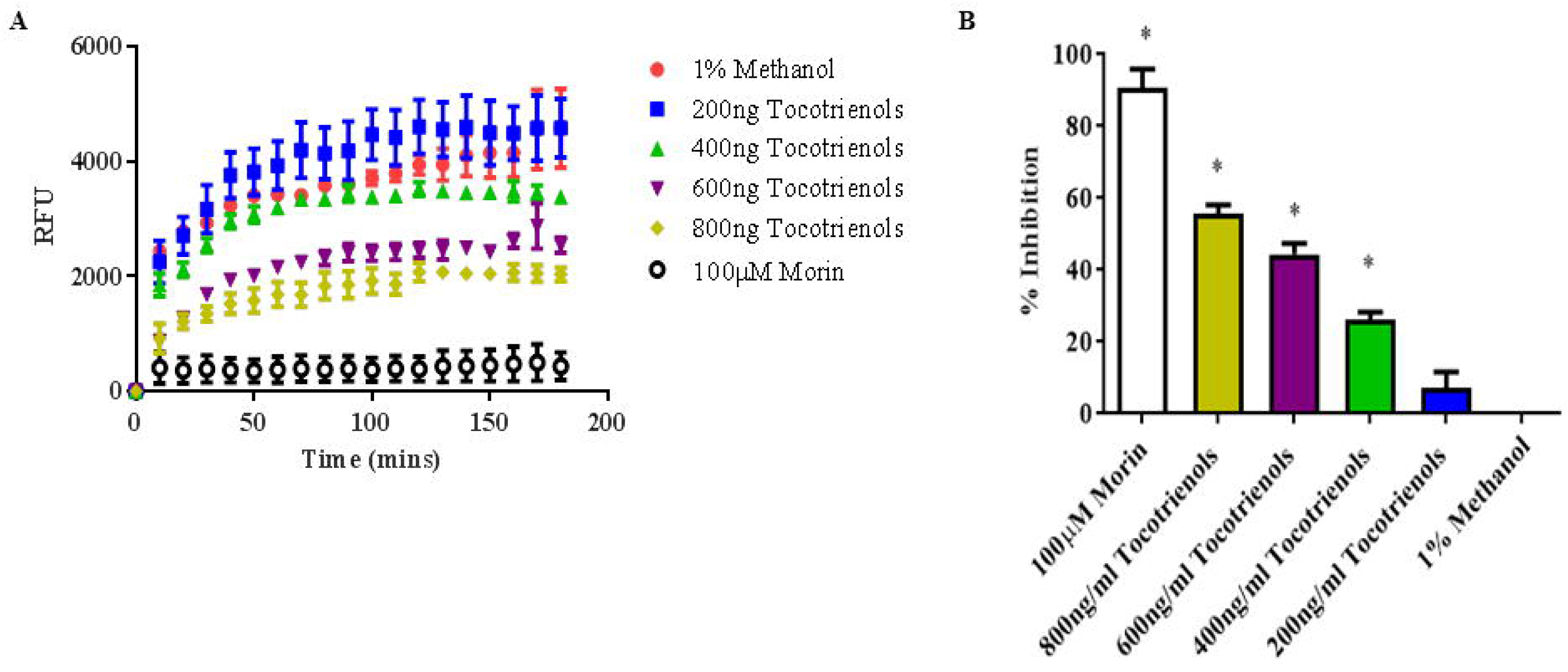
**A: Effects of mixed tocotrienols on Aβ aggregation.** Fluorescence signal was monitored at Ex/Em= 440/484 nm every 10 minutes at 37°C with 15 seconds between reads. l00µM morin acts as a positive inhibitor. **B: The percentage of inhibition of Aβ**_**42**_ **formation by various concentrations of mixed tocotrienols at 180 minutes.** Aβ_42_ aggregation was inhibited at concentrations of 400ng/ml, 600ng/ml and 800ng/ml (p<0.05) mixed tocotrienols. IC_50_ was 600ng/ml mixed tocotrienols.

Significant inhibition of Aβ fibrils formation was observed for 600 ng/ml and 800 ng/ml of mixed tocotrienols (p<0.05) compared to the two lower concentrations (Figure 2 A). The IC_50_ for the inhibition of Aβ aggregation by tocotrienols was 600 ng/ml (see Figure 2 B).

Tocotrienols are known to have antioxidant properties and were therefore tested using dichloro-dihydro-fluorescein diacetate (DCFH-DA) assay. This study showed the potential of mixed tocotrienols in reducing Aβ-induced ROS production (Figure 3 A). The worms were harvested at 32 hours after temperature upshift to observe the differences in ROS productions between the worms treated with mixed tocotrienols and the untreated. As shown in Figure 3 B, tocotrienols mixture demonstrated the potential in reducing the ROS production but did not totally negate it. Worms treated with 0.5 %-1 % mixed tocotrienols showed significantly reduced ROS production. The results were in correlation with the anti-paralysis assays. Based on Figures 4 and 5, 1 % mixed tocotrienols (equivalent to 100μg/ml mixed tocotrienols) affected the Aβ_42_ gene expression compared to methanol (used as the vehicle) but did not statistically significant decrease the Aβ_42_ levels (p>0.05).

**Figure 3.**
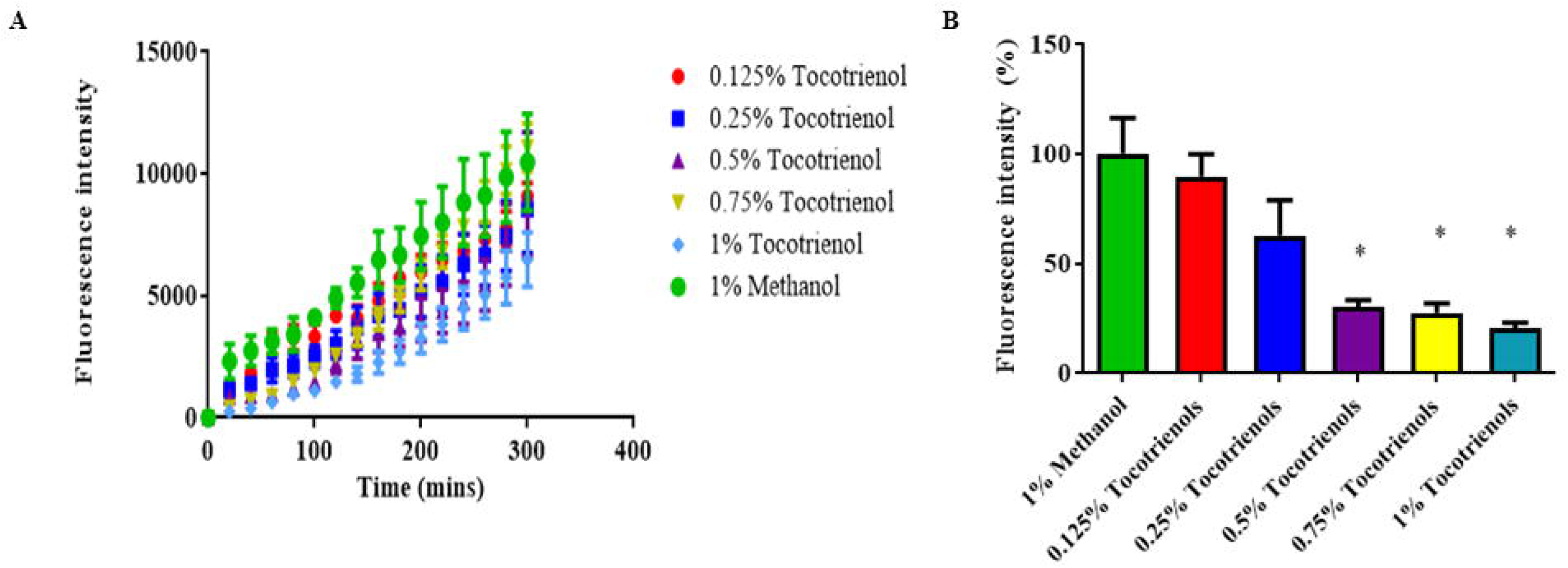
**A: Effects of mixed tocotrienols on the reduction of oxidative stress.** Comparison of relative oxygen species levels in transgenic *C. elegans* GMC101 treated with concentrations 0.125% to 1% mixed tocotrienols with the duration of 300 minutes with every 20 minutes readings. **B: Comparison relative oxygen species levels in transgenic *C. elegans* GMC101 treated with concentrations 0.125% to 1% mixed tocotrienols at the 60 minutes.** Oxidative stress in worms was significantly reduced at 0.5%, 0.75% and 1% mixed tocotrienols (p<0.05).

**Figure 4:**
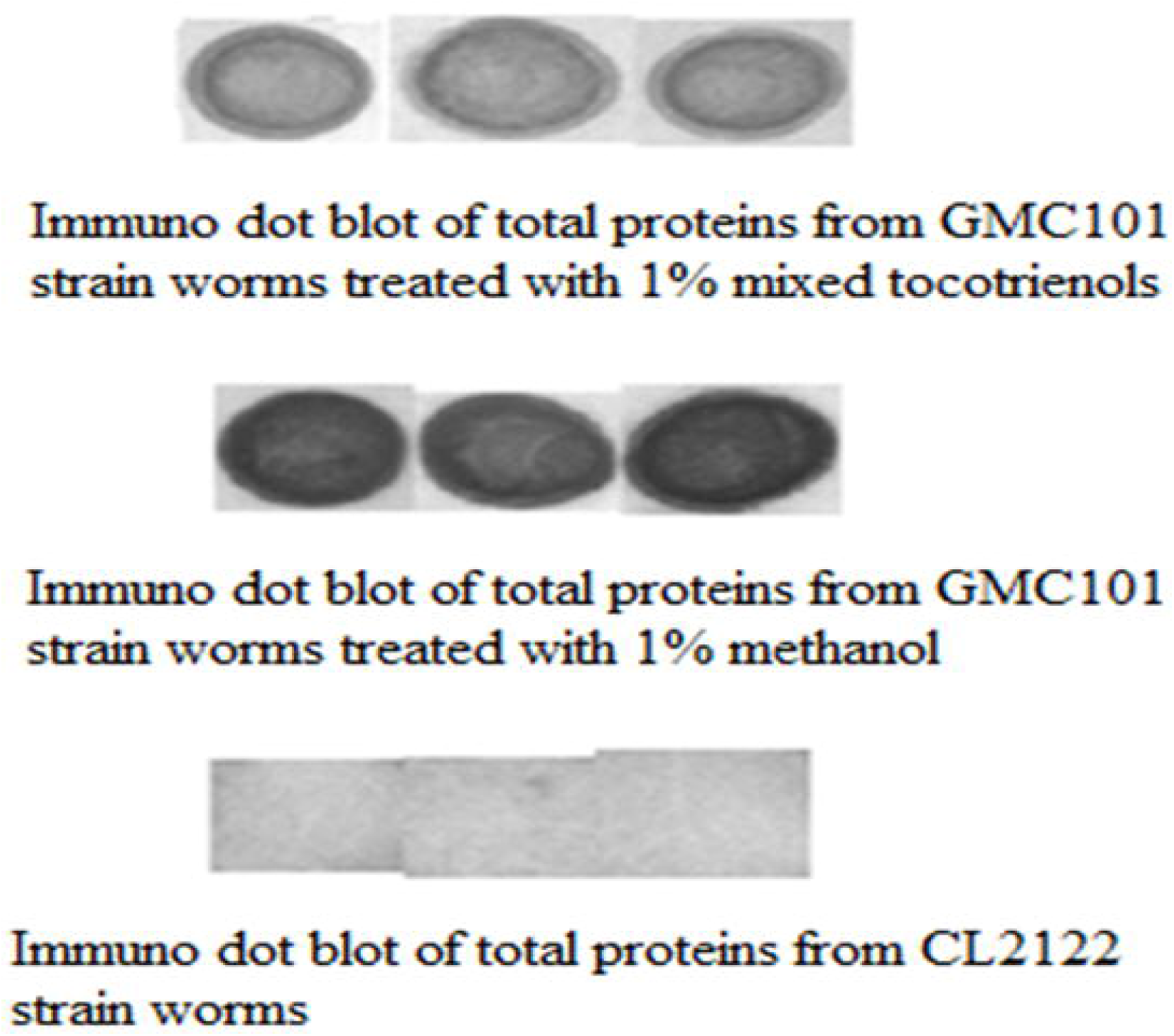
Effects of mixed tocotrienols on the Aβ levels. Representative dot blot of total Aβ in GMC101 transgenic worms fed with or without 1% mixed tocotrienols. No unspecific binding of Aβ antibody observed when total proteins of CL2122 were used.

**Figure 5:**
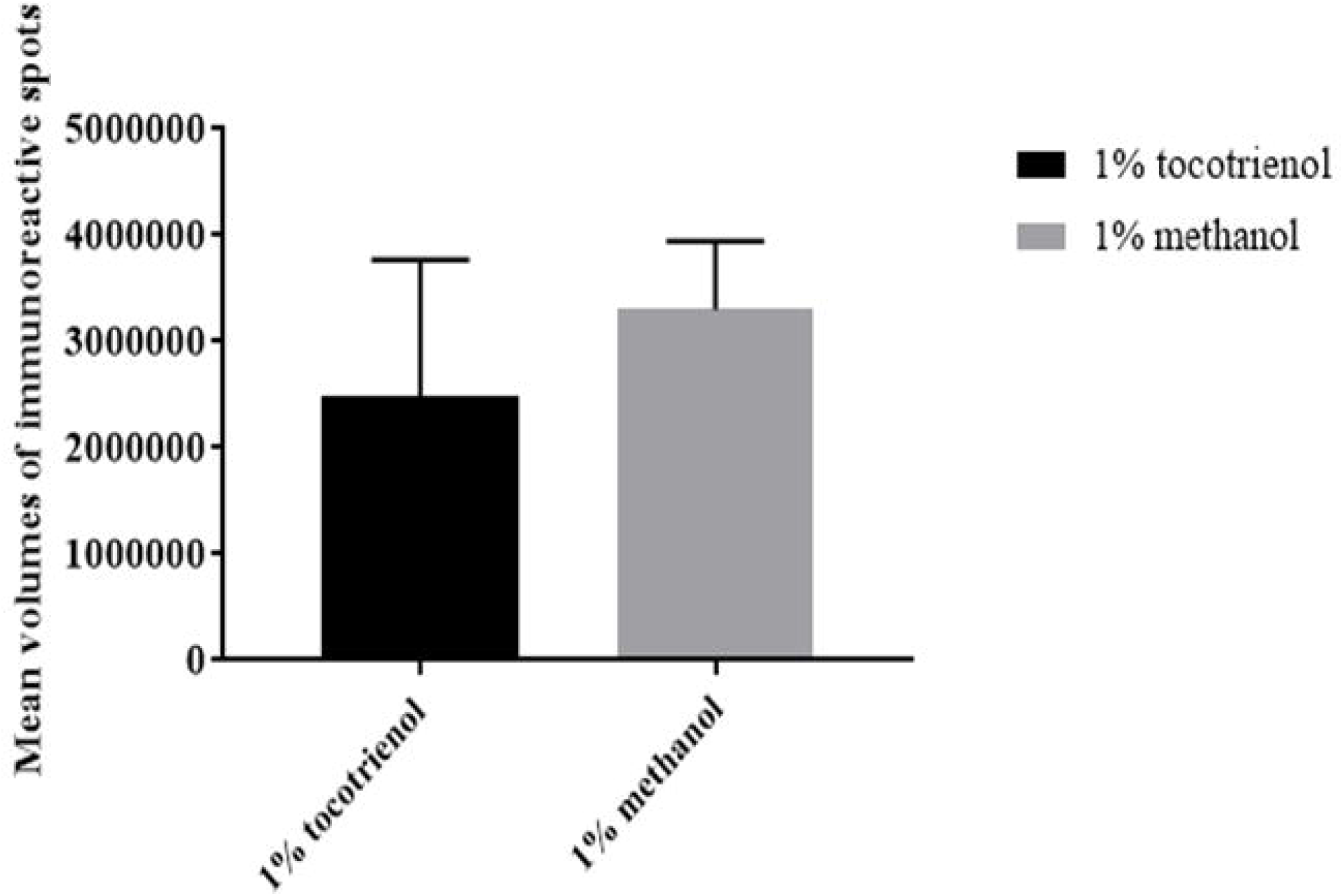
Exposure to mixed 1% tocotrienols affected the Aβ42 gene expression compared to methanol for 32 hours but did not significantly decrease the Aβ42 levels (p>0.05)

## 4.0 Discussion

A palm oil extract comprising mixed tocotrienols with a combination of 12.1% α-, 2.7% β-, 18.6% γ- and 8.1% δ-tocotrienols was evaluated for its neuroprotection activities. Based on the paralysis assays on a *C. elegans* transgenic strain that expressed Aβ proteins, mixed tocotrienols delayed amyloid-induced paralysis. A study had shown that mixed tocotrienols with the combinations of 19.6% α-, 2.4% β-, 25.5% γ- and 7.5% δ-tocotrienols had disrupted the Aβ oligomers and Aβ fibrils from being formed (5). In addition, Aβ depositions and fibrillar type plaques were reduced in the brains of mixed tocotrienols treated AβPPswe/PS1dE9 double transgenic mice (5). Cognitive function of the treated mice had also improved in the object recognition test as compared to control AβPPswe/PS1dE9 mice (5). Hence, it can be concluded that mixed tocotrienols had decreased amyloid pathology and improved cognitive functions.

For the *in vitro* study, the tocotrienols act directly towards the Aβ_42_ proteins and therefore the concentrations of tocotrienols used was lower than the concentrations used in the tocotrienol studies on the nematodes. The concentrations used on the worms were higher considering that the entry route was by ingestion and subsequent metabolism of the compounds would occur.

Mixed tocotrienols with combinations of 10.8% α-, 22.0% γ- and 5.2% δ-tocotrienols was shown to prevent hydrogen peroxide-induced death in rat striatal neuron cells. The minimal concentration that showed anti-H_2_O_2_-induced neurotoxicity properties for α-, γ- and δ-tocotrienols were 0.1, 1 and 10 μmol/L, respectively (9). The lowest dosage of mixed tocotrienols that posed anti-paralysis effects to the transgenic GMC101 strain was 0.5% mixed tocotrienols which was equivalent to 50 μg/ml. Thus, it can be deduced that tocotrienols have antioxidant properties.

In another study, protein oxidation was reduced in 15-day old *C. elegans* that was treated with mixed tocotrienols with the contents of 24% α-, 37% γ- and 12% δ-tocotrienols (10). In addition, protein oxidation was effectively inhibited in skeletal muscles of rats when rats were treated with tocotrienols supplements (11). In the present work, mixed tocotrienols with 12.1% α-, 2.7% β-, 18.6% γ-, and 8.1% δ-tocotrienols had reduced Aβ_42_-induced oxidative stress in the transgenic *C. elegans*.

## 5.0 Conclusion

In this study, we found that tocotrienol treatment towards *C. elegans* expressing human Aβ_42_ gene showed positive response. From the analysis, paralysis was delayed when *C. elegans* was treated with tocotrienol. In addition, tocotrienol also inhibits the formation of Aβ fibrils as well as reducing Aβ-induced ROS production in *C. elegans*. Through this study, we infer that tocotrienol has the potential to be an alternative drug to combat Alzheimer’s disease.

## Acknowledgement

We would like to thank all our collaborators and colleagues for the discussion and the work conducted in this lab. This study was funded by the USM Top Down Research Fund – URICAS (1001/PBIOLOGI/870029)

## References

1. Sadigh-Eteghad S, Sabermarouf B, Majdi A, Talebi M, Farhoudi M, Mahmoudi J (2015) Amyloid-beta: a crucial factor in Alzheimer’s disease. Medical Principles and Practice 24 (1):1–10 doi: 10.1159/000369101

2. Selkoe DJ (1994) Cell biology of the amyloid beta-protein precursor and the mechanism of Alzheimer’s disease. Annual review of cell biology 10 (1):373–403 doi: 10.1146/annurev.cb.10.110194.002105

3. Braak H, Braak E (1991) Demonstration of amyloid deposits and neurofibrillary changes in whole brain sections. Brain pathology 1 (3):213–216 doi: 10.1111/j.1750-3639.1991.tb00661.x

4. Sen CK, Khanna S, Rink C, Roy S (2007) Tocotrienols: the emerging face of natural vitamin E. Vitamins & Hormones 76:203–261 doi: 10.1016/S0083-6729(07)76008-9

5. Ibrahim NF, Yanagisawa D, Durani LW, Hamezah HS, Damanhuri HA, Wan Ngah WZ, Tsuji M, Kiuchi Y, Ono K, Tooyama I (2017) Tocotrienol-rich fraction modulates amyloid pathology and improves cognitive function in AβPP/PS1 mice. Journal of Alzheimer’s Disease 55 (2):597–612 doi: 10.3233/JAD-160685

6. Link CD (1995) Expression of human beta-amyloid peptide in transgenic Caenorhabditis elegans. Proceedings of the National Academy of Sciences 92 (20):9368–9372 doi: 10.1073/pnas.92.20.9368

7. Gutierrez-Zepeda A, Santell R, Wu Z, Brown M, Wu Y, Khan I, Link CD, Zhao B, Luo Y (2005) Soy isoflavone glycitein protects against beta amyloid-induced toxicity and oxidative stress in transgenic Caenorhabditis elegans. BMC neuroscience 6 (1):54 doi: 10.1186/1471-2202-6-54

8. McColl G, Roberts BR, Pukala TL, Kenche VB, Roberts CM, Link CD, Ryan TM, Masters CL, Barnham KJ, Bush AI (2012) Utility of an improved model of amyloid-beta (Aβ 1-42) toxicity in Caenorhabditis elegans for drug screening for Alzheimer’s disease. Molecular neurodegeneration 7 (1):57 doi: 10.1186/1750-1326-7-57

9. Osakada F, Hashino A, Kume T, Katsuki H, Kaneko S, Akaike A (2004) α-Tocotrienol provides the most potent neuroprotection among vitamin E analogs on cultured striatal neurons. Neuropharmacology 47 (6):904–915 doi: 10.1016/j.neuropharm.2004.06.029

10. Adachi H, Ishii N (2000) Effects of tocotrienols on life span and protein carbonylation in Caenorhabditis elegans. The Journals of Gerontology Series A: Biological Sciences and Medical Sciences 55 (6):B280–B285 doi: 10.1093/gerona/55.6.b280

11. Chung E, Mo H, Wang S, Zu Y, Elfakhani M, Rios SR, Chyu M-C, Yang R-S, Shen C-L (2018) Potential roles of vitamin E in age-related changes in skeletal muscle health. Nutrition Research 49:23–36 doi: 10.1016/j.nutres.2017.09.005

